# Context-dependent benefits of training and reminders in visual skill learning

**DOI:** 10.1101/2025.11.14.688438

**Authors:** Ylka Kolken, Mark J. Roberts, Parisa Naseri, Federico De Martino, Nitzan Censor, Peter De Weerd

## Abstract

Previous studies using a visual texture discrimination task (TDT) have demonstrated that performance enhancements resulting from extensive daily training (*full training* condition) remained intact after replacing all training, except for the first and last session, with a few daily reminder trials (*reminder* condition). Omitting reminders (*control* condition) yielded only limited learning, supporting their crucial contribution. We first confirmed these findings and excluded gaze position differences among conditions as a contributing factor. Next, we tested whether the reminders’ effectiveness is specific to a context of limited attention to the peripheral target caused by simultaneously performing a demanding fixation task. Removing the fixation task yielded performance levels in the first session matching those normally reached after lengthy daily training, suggesting that learning in the standard TDT involves the redeployment of attention. After changing texture parameters to increase the difficulty of the task, performing the TDT without a fixation task yielded learning in all three conditions. This indicates that in a dual-task, reminders can produce learning outcomes comparable to *full training*. In contrast, when the TDT is performed with full attention to the target, consolidation of the initial session alone can yield improvements equivalent to those observed in *reminder* and *full training* conditions.

## Introduction

Visual Perceptual Learning (VPL) refers to the enhancement of perceptual abilities through experience and training. As early as the nineteenth century, James (1890) and Stratton (1896), emphasized the importance of extensive daily practice in acquiring habits and refining skills. Karni and Sagi (1991) aptly captured this view in one of their hallmark papers, entitled “Where Practice Makes Perfect […]”. Numerous studies have supported the idea that visual skills develop over weeks or months of deliberate practice (Gibson and Gibson, 1955; McKee and Westheimer, 1978; Karni and Sagi, 1991; Fahle and Edelman, 1993; Schoups et al., 1995; Seitz et al., 2005; Yotsumoto et al., 2008; Gilbert et al., 2009; Lu et al., 2016; Watanabe and Sasaki, 2015). Contrary to this view, a recent series of studies has demonstrated that brief memory reminders produce performance gains comparable to those achieved through daily training, questioning the necessity of intensive, daily practice (Amar-Halpert et al., 2017; Kondat et al., 2023, 2024). Using a texture discrimination task (TDT), these studies demonstrated that after an initial training session, only five daily near-threshold trials (reminders) on subsequent days were sufficient to match performance gains seen with daily practice. Amar-Halpert et al. (2017) proposed that reminder-induced reconsolidation cycles (Walker et al., 2003; Lee, 2008; Dudai, 2012; Nader, 2015) enable efficient perceptual learning, making extensive daily training redundant. Here, we investigated the context in which reminders enable learning.

The TDT paradigm used by Amar-Halpert et al. (2017) and others (Karni and Sagi, 1993; Stickgold et al., 2000; Harris et al., 2012; Meng et al., 2014; Kondat et al., 2023, 2024) was originally devised by Karni and Sagi (1991). It is a dual task (from now on referred to as’dual-task TDT’) in which participants fixate and discriminate centrally presented T/L targets while judging the orientation (horizontal vs. vertical) of a peripheral texture-defined target. Task difficulty of the dual-task TDT is manipulated via the Stimulus Onset Asynchrony (SOA) between the texture stimulus and a following mask, and performance is measured as the SOA yielding 81.6% accuracy. The T/L task is included to ensure fixation without the need for eye-tracking equipment. Because the dual-task TDT requires simultaneous central and peripheral processing, its difficulty may partly stem from dividing attention between two relevant locations. The idea that dividing attention in dual task conditions incurs a cost for one or both tasks is well-documented (for a review see (Jans et al., 2010)). Accordingly, Kastner et al. (1998) showed that when demanding central T/L and peripheral target discrimination tasks were performed simultaneously, peripheral performance dropped from 86% to 45%. In the context of VPL, studies that have explicitly manipulated attention, have reported reduced learning under divided compared to full attention (Mukai et al., 2011; Meng et al., 2014). We therefore hypothesized that reminder benefits as studied in Amar-Halpert et al. (2017) are enabled by suboptimal learning due to limited attentional resources. In other words, reminders may only introduce benefits when previous learning was poor due to the dual task.

An alternative account for learning in the dual-task TDT, and potentially for differences in learning among conditions, is related to anticipatory gaze positioning prior to stimulus presentation. We reasoned that training-induced performance enhancements in the peripheral task with training may result from increased allocation of attentional resources to the peripheral target, as more resources are freed when participants improve in the central task. Because attention and eye movements are closely linked (Hoffman, 1998; Schroth et al., 2025), participants learning the task may unwittingly fixate their gaze closer to the anticipated peripheral target location, adopting a ‘compromise location’ to optimize performance in both tasks. This possibility so far has not been tested, because the T/L task at fixation has been considered a sufficient guarantee for accurate fixation, rendering the monitoring of eye position unnecessary. We nevertheless addressed this possibility by recording participants’ eye position during the dual-task TDT.

In summary, to examine whether reminder effectiveness depends on attentional division, we compared learning in the standard TDT (’dual-task TDT’) with learning in a version omitting the fixation task (’single-task TDT’). In both versions of the TDT, participants were assigned to one of three conditions: training over five consecutive days (*full training* condition); training only on day 1 and 5 with five near-threshold reminders on days 2-4 (*reminder* condition); training only on day 1 and 5 without reminders (*control* condition). We predicted that in the dual-task TDT, the *full training* and *reminder* condition would show similarly substantial learning, whereas the *control* condition would show limited learning. In the single-task TDT, without a demanding fixation task, we expected that reminders would lose their effectiveness, and that only the *full training* condition would produce substantial performance improvements.

## Methods and Materials

### Participants

Ninety-five volunteers naive to the purpose of the experiment participated in the different experiments (for the distribution of participants among experiments, see Supplementary Table S1). Each experiment included three conditions (Figure 1): a *full training* condition, in which participants trained for five consecutive days; a *reminder* condition, in which participants trained on day 1 and day 5 and received five near-threshold reminder trials on days 2-4; and a *control* condition in which no training occurred on days 2-4. For all participants, the interval between the first (Day 1) and the final session (Day 5) was exactly 4 days. Participants were not informed about the experimental design or aims. Group assignment was disclosed only after the first training session. This ensured that all participants began the experiment with equal motivation and without bias from prior knowledge of their training schedule.

**Figure 1:**
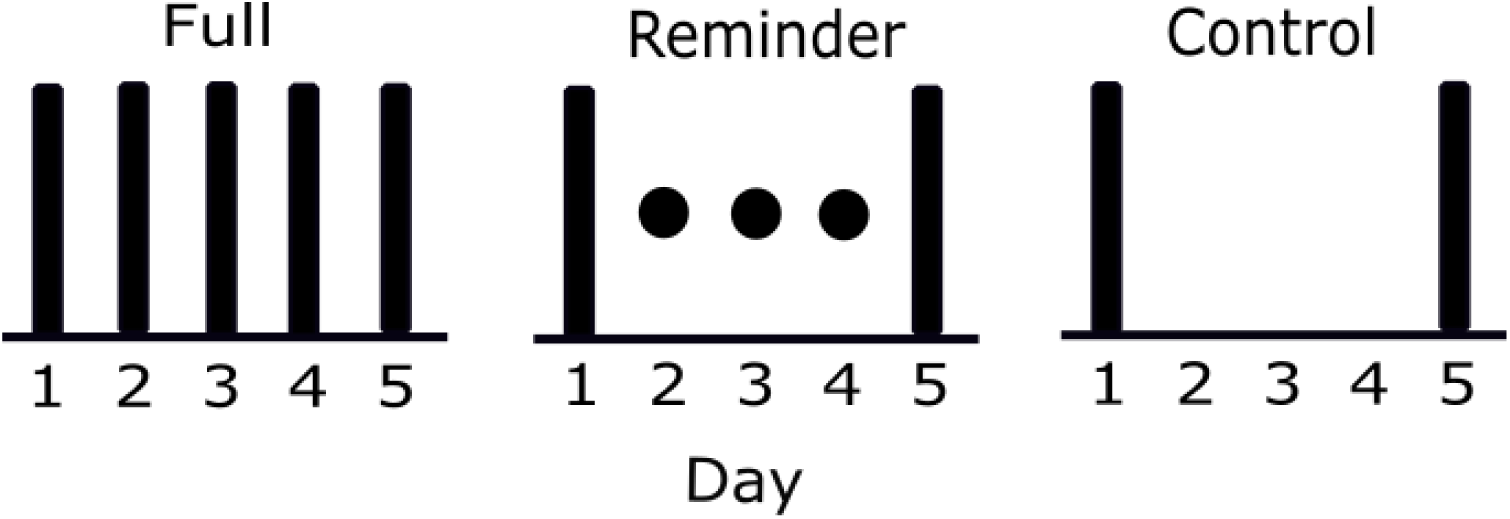
The three conditions used in all experiments; *full training*, *reminder*, and *control*. Stripes indicate a full training session of 252 trials. Dots indicate reminder sessions of 5 trials.

### Stimuli and tasks

We adopted the setup of Amar-Halpert et al. (2017) with the only difference being a longer stimulus duration (17ms instead of 10ms) due to the lower refresh rate of our screens. Participants discriminated the orientation of a texture-defined target stimulus presented in the lower-right visual quadrant (Figure 2A). The target stimulus consisted of three diagonal bars, presented in the lower right quadrant of the visual field at 5.72° eccentricity (x = 4.042°, y=-4.042°, Figure 2B). During transfer testing on Day 5, the texture-defined target appeared in the upper-left quadrant at the same eccentricity (x =-4.042°, y = 4.041°). The background array comprised 19 × 19 bar elements (0.57° X 0.04°), spaced 0.86° apart with 0.04° jitter. The display subtended 15.4° X 15.1° (24-inch Iiyama Prolite PC monitor, 60Hz refresh rate, 1920 x 1080-pixel resolution, mean texture luminance 84 cd/m2), viewed from 65cm distance. All target durations, SOAs, and the mask duration were verified with a photodiode.

**Figure 2:**
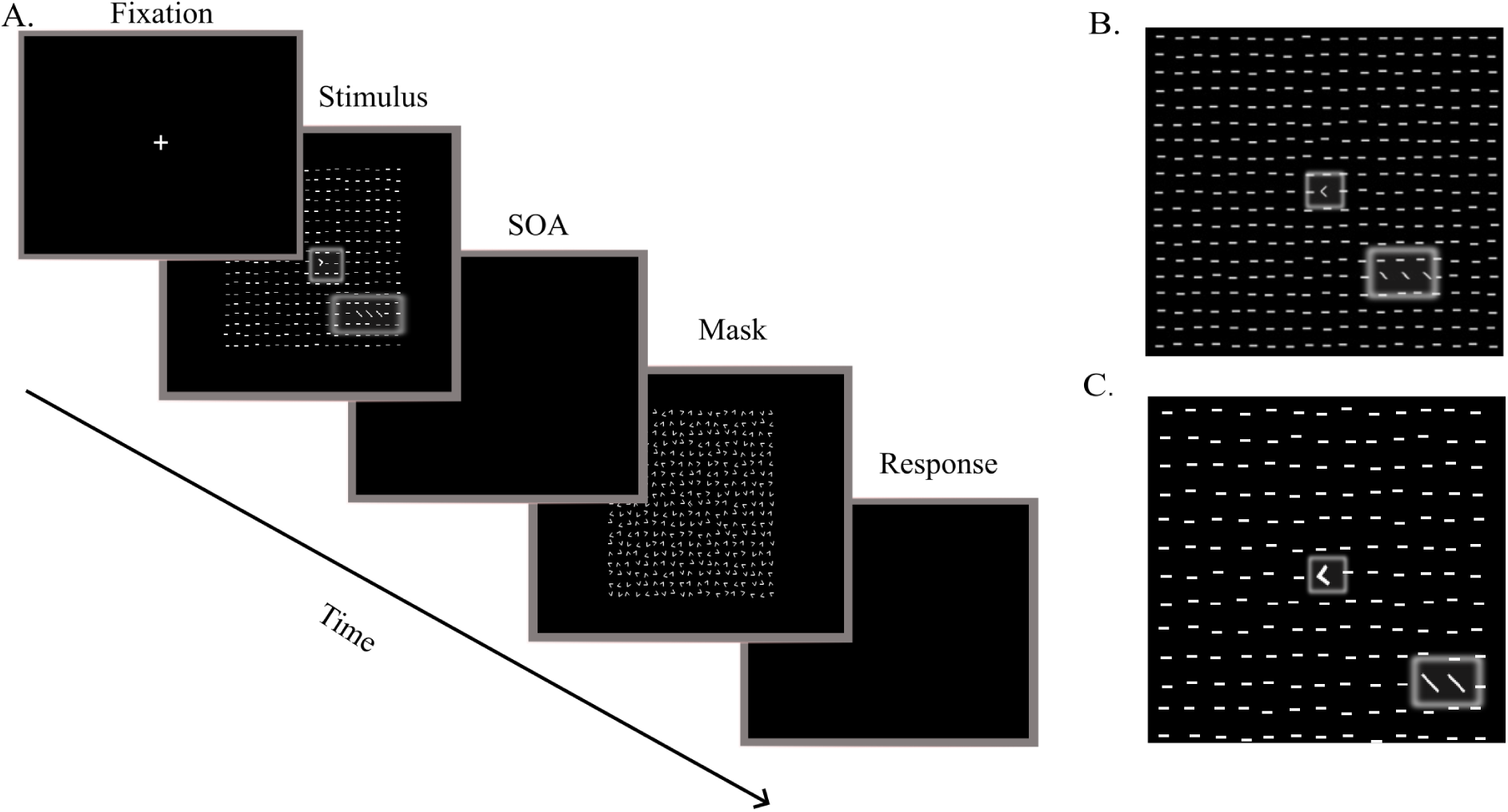
A. Trial sequence showing successive stimulus and mask presentation separated by the stimulus onset asynchrony (SOA). The white boxes inserted in the target frame indicate the location of T/L at fixation, and the texture target in the periphery. These boxes are for illustration and were not shown to the participant. Participants used key presses to respond to fixation and peripheral targets, respectively. Auditory feedback was given following incorrectly responding to a fixation target; no feedback was provided after responding to the peripheral texture target. B. Stimulus used in the dual-task TDT (enlarged from the stimulus frame in A). C. Modified stimulus (wider texture spacing and two target elements instead of three) used to increase difficulty in the single-task TDT. The central T or L character remained present in the texture, but participants were asked only to discriminate the orientation of the peripheral texture target.

In the dual-task TDT, fixation was facilitated by a forced-choice letter discrimination task (T or L at the centre of the display) with auditory feedback, in line with previous studies using this paradigm (Karni and Sagi, 1991; Yotsumoto et al., 2008; Kondat et al., 2023). Eye movements were recorded using a Viewpoint Eye Tracker (v2.8.3, Arrington Research). At the beginning of each trial, participants fixated their gaze on the center of the screen and pressed the space bar to start the trial (Figure 2A). After the appearance of a fixation cross, a target frame (17ms) was followed (after a given SOA) by a patterned mask consisting of randomly oriented V-shaped elements and a central composite pattern of superimposed T and L shapes (102ms). After the stimulus presentation, participants reported the perceived letter and target orientation with separate keyboard presses; first they reported the T or L fixation target using ‘T’ or ‘L’ keys, which was followed by auditory feedback in the case of incorrect response. Second, they reported the target orientation, using ‘T’ for ‘horizontal’ and ‘L’ for ‘vertical’, which was not followed by feedback. There was no response time limit.

Prior to the experiment, participants underwent a pre-training phase followed by a familiarization phase. In the pre-training phase, participants were given a maximum of ten blocks, each containing ten trials, with the aim of reaching a level of 90% correct responses in a block at the highest SOA (323ms) for both the fixation and the texture discrimination tasks. If the participant did not reach this level of accuracy within 10 blocks, the participant was dismissed. The familiarization phase consisted of a block of trials, presenting one trial per SOA. The intervals between the target and the mask onset (stimulus onset asynchrony, SOA), ranged from 17ms to 323ms in increments of 17ms and were randomized across all trials [17 51 68 85 102 119 153 170 187 204 221 255 289 and 323]. Participants who did not reach the 81.6% correct threshold after pre-training on day 1 were excluded from further participation. This criterion led to the exclusion of 15 participants. Of the 30 included participants, three in *full training* were trained with 13 SOA levels instead of 14 (S3 on all days, S12 only on day 1, S5 on day 3) due to a video card error, and one participant in *full training* performed only 7 blocks on day 1 instead of 9 due to time pressure (S14).

In the single-task TDT, we used the same setup as in the dual-task TDT, except that the T/L task was omitted. In the absence of a central fixation task, fixation was enforced using an eyetracker (Arrington Research, Scottsdale, AZ, USA). If the participants’ gaze deviated *>* 1.5*^◦^* away from the center of the screen at any point during a one-second interval prior to stimulus presentation, or during stimulus presentation, the trial was aborted and replaced by another trial. The aborted trial was re-administered later in the session. An automatic recalibration of the eye tracker would begin if participants broke fixation on 5 consecutive trials.

Ten participants performed the single-task TDT with the same stimuli as used in the dual-task TDT in a single session. The single-task TDT was also performed with a modified stimulus, which consisted of a 13 × 13 element texture array (line elements of 0.57 × 0.04°, spaced 1.2° apart with 0.05° jitter; see Figure 2C), instead of 19 × 19 array, and 2-element targets instead of 3-element targets. These changes were motivated by preliminary data, which showed near-maximum performance could be reached in the first session with the original stimulus (see Supplementary section 3). With this modified stimulus, we trained participants in the same three conditions (*full training*, *reminder*, and *control*) as in our dual-task TDT.

### Sample size

To determine the required sample size for the dual-task TDT, we based our estimation on the a priori difference between the *full training* and *control* conditions, as a sufficiently robust difference between *control* and *full training* conditions is a prerequisite for a meaningful comparison of the *reminder* condition with these two conditions. In the study by Amar-Halpert et al. (2017), the mean relative improvement in perceptual thresholds from day 1 to day 5 was 26.6% for the full-training condition (N = 12) and 6.3% for the control condition (N = 7) (assuming similar Day 1 thresholds, which were not reported). Based on the reported group-level variability, we estimated a pooled standard deviation of 17%, yielding a corresponding Cohen’s d *≈* 1.17 for the group difference. An a priori power analysis conducted in G*Power (Version 3.1; Erdfelder et al. (1996)) indicated that detecting this effect size with 80% power (1-*β*) at *α* = 0.05 would require 10 participants per condition, which we therefore adopted in our design. For the single-task TDT, we used the same effect-size estimate and applied the same rationale, and likewise planned for 10 participants per condition.

### Threshold calculation

The psychometric function was modeled using the Weibull sigmoid, scaled to account for guessing and lapses, following the form used in the psignifit4 toolbox (Wichmann and Hill, 2001):

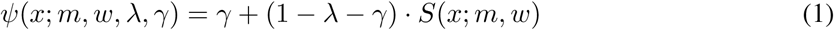

where threshold *m* is the SOA level at which 81.6% of responses are correct; *w* represents the width; *λ* represents the lapse rate (free); and *γ* represents the guess rate, fixed at 0.5.

### Statistics

Frequentist analyses were performed in Python 3.10 (Van Rossum and Drake, 2009), Bayes factor calculations in JASP (JASP Team, 2025), and eye-tracking analyses in MATLAB (The MathWorks Inc., 2024). To check for baseline differences, we first conducted a one-way ANOVA on the Day 1 thresholds to test for condition differences. To evaluate both within-condition learning and between-condition differences in learning across the *full training*, *reminder*, and *control* conditions, we fit a linear mixed-effects model (LMM). This model included fixed effects of Time (Day 1 vs. Day 5), Condition, and their interaction (Time × Condition), allowing us to assess overall changes in performance, condition differences in baseline performance, and whether learning (i.e., performance change over time) varied by condition.

Random intercepts and slopes for Time were included for each subject to account for individual differences in initial performance and learning rate. The model was specified as follows:

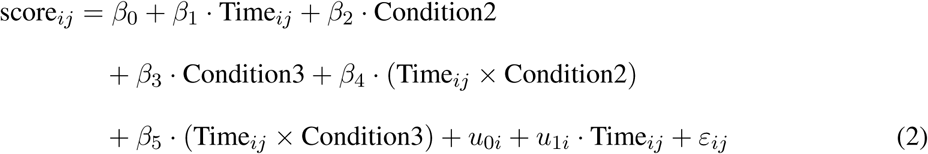

where score*_ij_* and time*_ij_* represent respectively the performance score (in ms) and the measurement time for the *j*th measurement of the *i*th participant. *Condition* stands for the three conditions (*control* is condition 1 (baseline), *full training* is condition 2, and *reminder* is condition 3). The fixed effects are represented by *β* coefficients: *β*_0_ is the overall intercept (i.e., the grand mean performance), *β*_1_ captures the average effect of Time (change from Day 1 to Day 5), *β*_2_ and *β*_3_ represent the effect of Condition, and *β*_4_ and *β*_5_ capture the interaction between Time and Condition, which reflects differences in learning across conditions. The terms *u*_0_*_i_* and *u*_1_*_i_* are subject-specific random effects: *u*_0_*_i_* is a random intercept accounting for individual differences in baseline performance, and *u*_1_*_i_* is a random slope allowing for individual variability in learning rate (i.e., change over time). The residual error term *ε_ij_* is a within-individual measurement error that captures unexplained variance at each observation. Together, these components allow the model to flexibly account for both fixed effects of experimental factors and random variation across individuals. Model assumptions were evaluated by inspecting residuals against fitted values to ensure homoscedasticity and normally distributed errors.

In addition to the frequentist mixed-effects model, we conducted a Bayesian ANOVA on normalized Day 5 scores (calculated as Day 5 performance minus Day 1 performance) to assess condition differences in learning. This analysis was performed using JASP (version 0.19) with default priors: a zero-centered Cauchy 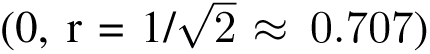 for fixed effects and a Jeffreys prior for variance components which lead to non-informative priors. The Bayesian approach permits quantifying the relative evidence for or against the presence of condition differences by computing Bayes factors BF, offering a complementary interpretation to traditional null hypothesis significance testing. The magnitude of the Bayes Factor reflects the strength of evidence (Jeffreys, 1961; Rouder et al., 2012; van den Bergh et al., 2019). We report *BF*_10_, which quantifies the evidence in favor of the alternative hypothesis (H1) relative to the null hypothesis (H0). A *BF*_10_ *>* 1 indicates support for the alternative hypothesis, whereas A *BF*_10_ *<* 1 indicates support for the null hypothesis.

To quantify changes in gaze behavior over time, we analyzed the eye-tracking data collected on Day 1 and Day 5 of the experiment. For each participant in these two sessions, we computed the distance of the average gaze position from the center of the screen to assess potential shifts in overall gaze centering across sessions. Second, we calculated the distance between the average gaze position and the target location, to see if, over time, participants shifted their gaze towards the target. To evaluate whether these measures changed over time, we performed a paired samples t-tests comparing Day 1 and Day 5 across participants. We used circular statistics to test whether gaze shifts were consistent in direction across participants by submitting Day 1–Day 5 movement angles to a Rayleigh test of non-uniformity. Fixation behavior was stored only in the dual-task TDT and not in the single-task TDT, which restricted this analysis to the dual-task TDT experiment.

Location transfer within each condition was assessed using paired t-tests in the dual-task TDT. For the single-task TDT, too few participants completed the transfer test to allow meaningful condition analysis; therefore, all transfer analyses are reported in the Supplementary materials.

## Results

### Dual-task TDT

We collected 30 complete dual-task TDT datasets, divided equally over the *full training*, *reminder*, and *control* condition. The learning curves (Figure 3A) showed no threshold differences on Day 1 between participants in the three conditions (F(2,54)=0.094), p=0.910). Figure 3A shows a strong reduction in SOA in the *full training* and *reminder* conditions, but not the *control* condition. Accordingly, the LMM showed a significant Time x Condition interaction for *full training* versus *control* (*β*=-77.120, SE=31.27, p=0.014), *reminder* versus *control* (*β*=-67.827, SE=31.27, p=0.03), but not between *full training* and *reminder* (*β*=9.292, SE=31.27, p=0.766) (Figure 3A, for single-subject Day 1 versus Day 5 thresholds see Figure 3B). These results are in line with the reminder trials being as effective as *full training* (for all LMM effects and additional analyses, see Supplementary Section 2).

**Figure 3:**
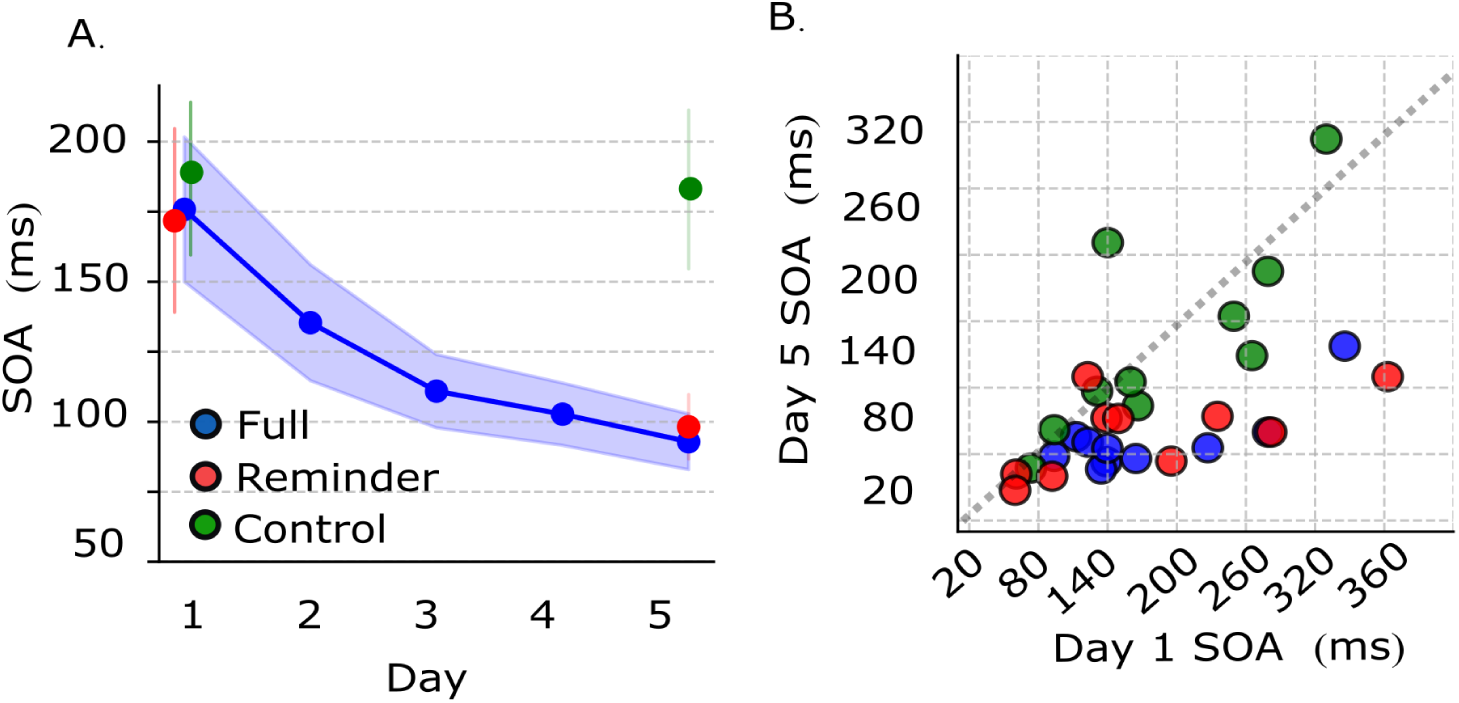
Comparison of learning outcomes between *full training* (in blue), *reminder* (in red), and *control* (in green) conditions in the dual-task TDT. A. Learning curves for the three conditions in the dual-task TDT. Error bars show 1 SEM. Day 1 thresholds are plotted with a small horizontal offset to display the SEM. B. Individual participant thresholds from day 5 plotted as a function of their day 1 thresholds. Data are colour coded as in panel. A. Dashed diagonal line corresponds to absence of learning, points falling below the diagonal indicate improved performance.

The Bayesian analysis confirmed that the *reminder* and *full training* conditions performed similarly (*BF*_10_ =.409), while the *control* condition differed from both the *reminder* (*BF*_10_ = 1.579) and the *full training* (*BF*_10_ = 5.036) conditions. This confirms our initial findings and shows that the *full training* and *reminder* conditions showed learning, whereas the *control* condition did not.

The learning results were not related to changes in fixation strategy. We had hypothesized that performance increases in the dual-task TDT were related to participants positioning their gaze closer to the anticipated position of the peripheral target. Our eye movement recordings do not support this idea. Overall, participants fixated close to the fixation spot, with 90% of all trials falling within a 2.4° radius from fixation on Day 1 (Figure 4A - left), and within a radius of 2.3° radius on Day 5 (Figure 4A - right). Moreover, there was no significant change in the average distance between the gaze point and the centre of the screen from day 1 to day 5 (t(27)=-0.038, p=.9699) or from the gaze point to the target from day 1 to day5 (t(27)=0.0101, p=.992). Furthermore, we found no consistency in the direction of shift in mean gaze position between days 1 and 5 (z = 1.87, p =.154) (Figure 4B). This suggests that participants neither fixated on the peripheral target nor adopted a gaze-shifting strategy to improve performance in the dual-task TDT.

**Figure 4:**
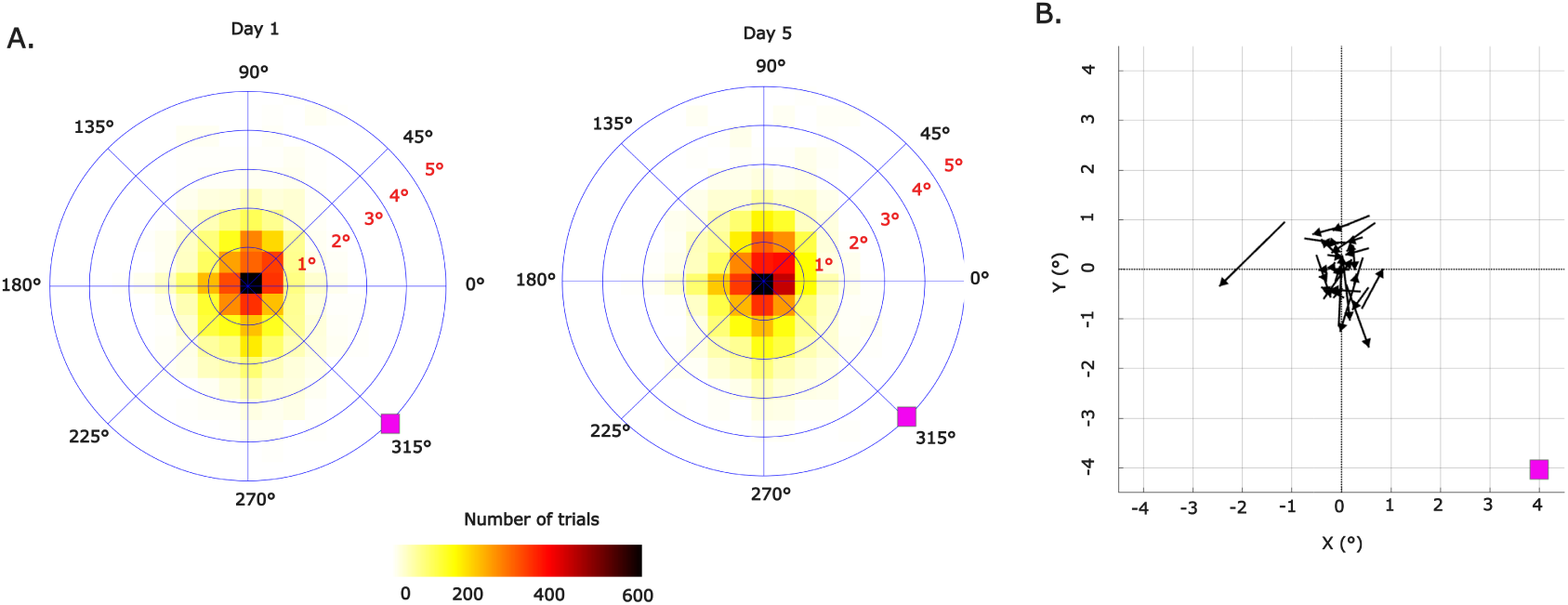
A. Fixation accuracy on day 1 (left) and on day 5 (right). B. Average X and Y positions per participant on day 1 (start of the arrow) and day 5 (arrowhead). Pink box denotes target position.

### Single-task TDT

In preliminary testing, 10 participants performed the TDT without the T/L fixation task using the same stimuli as the dual-task TDT. On the first day, their mean threshold was 62 ± 15.5 ms (median 53.58), much lower than the 180 ± 16.8 ms observed on Day 1 of the dual-task TDT, lower than any Day 5 thresholds across conditions, and lower than previously reported values (see Supplementary Figure S1 and Table S3). The low thresholds left little room for further improvement, so the experiment was aborted for these participants after Day 1. To increase task difficulty, we made the texture stimulus coarser while maintaining the dimensions of the texture elements. This manipulation is known to increase the difficulty of figure-ground texture segregation and hence the difficulty of the TDT task (Nothdurft, 1985; De Weerd et al., 1992)). Additionally, the figure was reduced to 2 elements (instead of 3), which further increased difficulty (for details, see Methods and Figure 2). Two additional changes were made relative to the dual-task TDT. First, the pre-training task was omitted, as it showed no correlation with day 1 performance (r = –0.217, p =.373) in the dual-task TDT and added no value beyond the familiarization task. Second, to reduce exclusions after modifying the stimulus on Day 1, the SOA range was adjusted: with the original range (17–323 ms), 8 of 15 participants failed to reach threshold, whereas with the modified range [51–425 ms, 13 levels], only 2 of 25 were excluded.

Using the single-task TDT with the modified stimulus, 30 participants completed the experiment (N=10 per condition). Performance on Day 1 showed no threshold differences between the three conditions (F(2,54)=0.018), p=0.98), and learning in the three conditions appeared similar (Figure 5A).

**Figure 5:**
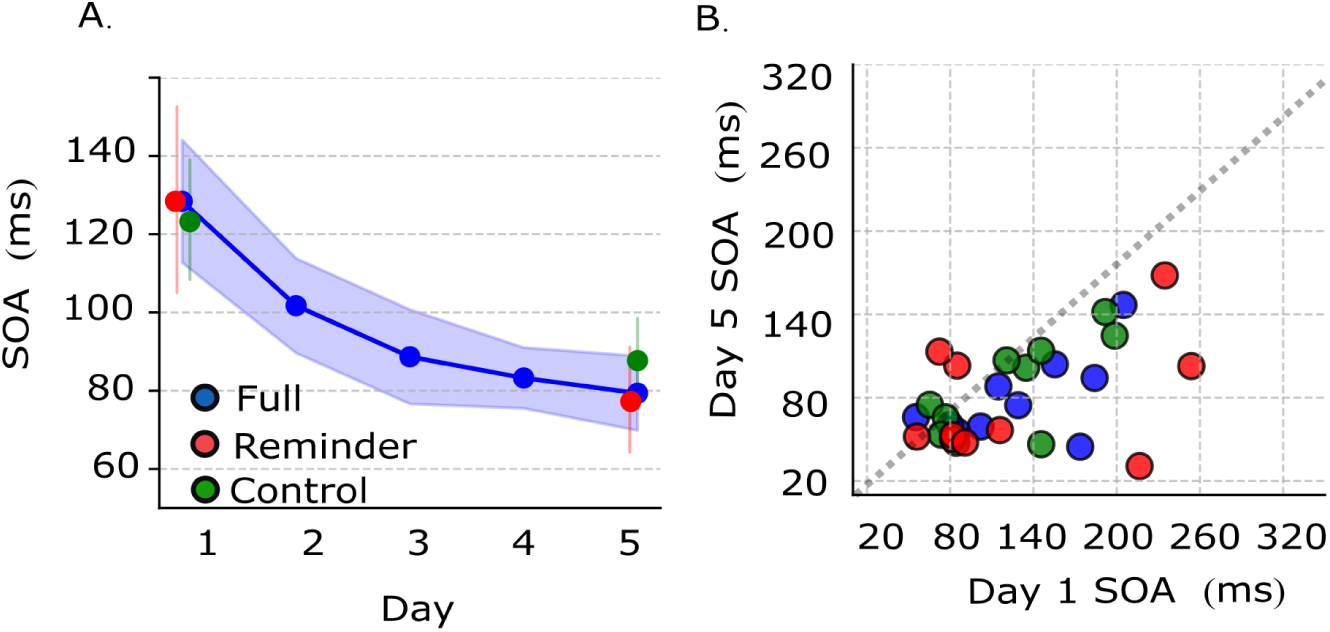
Comparison of learning outcomes among full training, reminder, and control conditions in the single-task TDT. All conventions as in Figure 3.

Accordingly, the LMM showed no significant Time x Condition interaction for *full training* versus *control* (*β*=-13.061, SE=20.561, p=0.525), *reminder* versus *control* (B=-8.169, SE=20.561, p=0.691), or between Full Training and *reminder* (*β*=4.892, SE=20.561, p=0.806) (Figure 5A, for single-subject Day 1 versus Day 5 thresholds see Figure 5B). After removing the interaction terms, the effect of Time effect was significant (*β*=-43.060, SE=8.161, *p <.*001). These results indicate that all three conditions showed a similar performance increase (for all LMM effects and additional analyses, see supplementary table S4 and S5). Bayesian analysis confirmed that the conditions performed similarly (*reminder* vs *full training* (*BF*_10_ =.404), *control* vs *reminder* (*BF*_10_ =.417) and *control* vs *full training* (*BF*_10_ =.505))

## Discussion

In a recent study, Amar-Halpert et al. (2017) demonstrated that visual skill leaning spanning five daily sessions (*full training* condition) could be matched by a training regimen in which several of the training sessions were replaced by just a few reminder trials (*reminder* condition). The reminder trials were crucial because omitting them (*control* condition) prevented learning. Here, we aimed to replicate these findings and to examine the mechanisms underlying reminder effectiveness. Because Amar-Halpert et al. (2017)) used a dual-task TDT, which constrains attention to the target, we hypothesized that the effectiveness of reminders is confined to learning conditions that limit attention to the target.

We successfully replicated Amar-Halpert et al. (2017)’s findings in the dual-task TDT. Unlike in their study, we equated the time difference between first and last session across all conditions, which renders the evidence for reminder effectiveness in our study arguably even stronger. Moreover, our eye-tracking data ruled out anticipatory gaze shifts towards the target as a source of improvement.

We next tested whether learning in the dual-task TDT (Karni and Sagi, 1991, 1993; Karni et al., 1994) occurs exclusively under conditions that limit attention to the target. Because participants in this task must simultaneously discriminate both a central target (T vs. L) and a peripheral target (horizontal versus vertical), we reasoned that attentional resources for the target would be constrained (Kastner et al., 1998; Jans et al., 2010). We hypothesized that these attentional constraints create the conditions under which reminders can be effective.

The difficulty of the experiment was evident in the high exclusion rates, with 15 of 45 participants failing to meet the pre-training or Day 1 performance criteria (see Methods). To test whether the difficulty in the dual-task TDT is attentional, we omitted the central fixation task, leaving all else unchanged. In this single-task version of the TDT, we observed low SOA thresholds on Day 1, comparable to thresholds in the dual-task TDT after five days of training (see Supplementary Figure S1). Thus, the high thresholds in the dual-task TDT on day 1 reflect attentional rather than sensory constraints. This likely applies to other studies that used the dual-task TDT (Karni and Sagi, 1991, 1993; Scialfa et al., 2000; Pavlovskaya et al., 2001; Amar-Halpert et al., 2017; Kondat et al., 2023), despite limited evidence to the contrary (Karni and Sagi, 1991).

To enlarge the possible learning amplitude in the single-task TDT, we modified the stimulus to reduce target discriminability (see Methods and Results). Hence, the enhanced difficulty in the single-task TDT was stimulus-related rather than attention-related. For the modified stimulus, we predicted *full training* would yield learning, whereas *reminder* and *control* conditions would not. Contrary to this hypothesis, all three groups—*full training*, *reminder*, and *control* — showed robust learning. Reminders had no additional benefit in the single-task TDT, but the unexpectedly strong learning in the *control* condition meant there was also no room for the reminders to outperform it.

The comparable learning in *control* and *full training* conditions suggests that the consolidation process following a single training session may last substantially longer than previously assumed, and is affected by neither reminder trials nor extensive daily training. Most skill learning studies employ daily training schedules (e.g. (Karni and Sagi, 1991; Schoups et al., 1995; Amar-Halpert et al., 2017), thereby implicitly assuming that consolidation does not extend beyond a single day. Similarly, interference studies (Brashers-Krug et al., 1996; Shadmehr and Brashers-Krug, 1997) have typically estimated consolidation durations to be in the order of a few hours, and sleep studies (e.g., (Fischer et al., 2002; Walker et al., 2002) have focused on overnight consolidation. Stickgold et al. (2000) however, demonstrated in a dual-task TDT that SOA thresholds decreased by approximately 20 ms when participants were retested three to four days after an initial training session, with no intervening sessions, supporting a protracted consolidation process. Our control data in the single-task TDT showed similar SOA reductions of about 36ms after five days without training. These results indicate that behaviorally relevant latent consolidation can extend over at least five days.

While we were conducting our study, Kondat et al. (2024) reported fMRI evidence implicating the intraparietal sulcus (IPS) in reminder-based learning, consistent with its known contributions to attentional control (Bisley and Goldberg, 2003), and to the division of attention in multi-target tasks (Prass and De Haan, 2017). Kondat et al. (2024) reported more activity in a rostral prefrontal region during *full training* than during *reminder*-based training. This region is associated with the default mode network and its activation could therefore be associated with a more automated mode of task execution. Kondat et al. (2024) thus suggested that *reminder*-based learning requires attentional contributions, whereas *full training* engage more automatic processes. Our data suggest a related but distinct interpretation. Because the dual-task context is present in all three conditions (*full training*, *reminder*, and *control*), we suggest attentional demands are required for all of them. Possibly, the differences in neural correlates between a full training block and a block of 5 reminder trials in Kondat et al. (2024) may reflect a stronger engagement of attention during the limited number of reminder trials than during the full block of training trials.

Amar-Halpert et al. (2017) attributed reminder benefits to reconsolidation – a process by which retrieved memories become labile and need to be reconsolidated (Nader et al., 2000; Nader, 2015). Reconsolidation can be disrupted by behavioral interference (Schroeder et al., 2023), stress (McGaugh, 2004; Cai et al., 2006), the inhibition of protein synthesis (Nader et al., 2000), or stimulation (Misanin et al., 1968; Robertson, 2012; Shmuel et al., 2021). Yet, the lability of the memory trace during reconsolidation may also allow strengthening through attention, consistent with related evidence that attention strengthens consolidation (Kentros et al., 2004).

Our and other’s findings (Amar-Halpert et al., 2017; Kondat et al., 2023, 2024) suggest that attentional state and stimulus design create the conditions under which reminders can be effective. This perspective may explain the recent non-replication of reminder-induced learning in a recent report by Zhu and Zhang (2025). They observed comparable learning in a dual-task TDT across *full training*, *control* and *reminder* conditions, mirroring our findings in the single-task TDT. Therefore, a potential explanation for Zhu and Zhang (2025)’s findings is that their participants had more attention available for the target than in preceding dual-task TDT studies. Differences in the size of the T/L stimuli at fixation, or differences among participant pools in pre-existing perceptual skills (e.g., due to gaming experience) and motivation (e.g., due to differences in instruction or in the perceived value of the reward) could all have modulated attentional allocation to the texture target in such a manner to mimic our single-task TDT results.

In summary, our data confirm that in an attention-demanding, 5-day dual-target visual perceptual learning task, reminders were beneficial in short-cutting daily training sessions. Learning did not rely on changes in fixation strategy. Furthermore, in a single-target task, permitting more attention to the target, training sessions following initial training could be entirely omitted without compromising performance in a final session. The reminder benefits obtained in dual tasks likely reflect reconsolidation processes, whereas the continued learning in single tasks without training for several days suggests a behaviorally relevant, protracted process of consolidation, after sufficient encoding of the memory on the first day of training. Future research should determine whether these effects extend to later, asymptotic learning phases.

