## Supplementary materials for "Context-dependent benefits of training and reminders in visual skill learning"

### 1 Participants

Table S1: Division of participants over the experiments

| Experiment | N included | N excluded |
|---|---|---|
| Dual-task TDT | 30 | 15 |
| Preliminary single-task TDT | 10 | 8 |
| single-task TDT 2 | 30 | 2 |

### 2 Dual TDT

#### Mixed Linear Model results

Table S2: Mixed Linear Model (dual-task TDT). Reference condition = Control, baseline = Day 1.

| Predictor | $\beta$ | SE | $z$ | $p$ |
|---|---|---|---|---|
| Intercept | 188.77 | 28.79 | 6.56 | $< .001$ |
| Time | -5.84 | 22.11 | -0.26 | .792 |
| condition 2 | -12.95 | 40.71 | -0.32 | .750 |
| condition 3 | -16.90 | 40.71 | -0.42 | .678 |
| Time × condition 2 | -77.12 | 31.27 | -2.47 | .014 |
| Time × condition 3 | -67.83 | 31.27 | -2.17 | .030 |

#### Supplementary results Dual-task TDT

To examine whether individual differences in reminder performance were related to overall learning, we tested for a correlation between participants' performance during the reminder sessions (defined as the percentage correct on all the reminder trials) and their overall learning (Day 5 - Day 1). This analysis was motivated by Herszage et al. (2023), who reported that the quality of reminder trials predicted subsequent learning gains in a motor skill task. No significant correlation was observed (r = -0.415, p = 0.2661), suggesting that splitting the reminder group based on reminder strength was not warranted.

In addition, to confirm that performance improvements in the *full training* condition reflected genuine learning rather than a test–retest effect, we compared thresholds between Day

2 and Day 5 within this condition. Participants improved significantly from Day 2 to Day 5 (t(9)=3.328, p=.0088), indicating that any results observed where not the results of a simple test-retest effect.

#### Transfer results dual-task TDT

We explored the extent of position generalization in the three conditions. We had speculated that in the *full training* condition, in which many more trials were presented than in the others, more specificity would emerge, either due to enhanced read-out or possibly due to low-level plasticity. Due to time constraints on Day 5, transfer thresholds were collected only in 26 out of 30 participants. Of those 26, 2 did not reach an 81.6% threshold in the secondary location (see Methods), leaving 24 participants for the analysis. In the *full training* condition, transfer thresholds were significantly higher than the day 5 thresholds at the trained location (t(8)=-3.95, p=.0042, cohen's d=-1.32), but not in the *reminder*(t(7)=-2.19, p=0.0647, cohen's d=-0.77) and *control* condition (t(6)=0.07, p=0.9480, cohen's d=0.03).

### 3 Single-task TDT

#### Preliminary testing

In preliminary testing, 10 participants performed the TDT without a fixation task using the same stimuli as in the dual TDT. On the first day, their mean threshold was 62 ± 15.5 ms (median 53.58; leftmost bar in S1), much lower than the 180 ± 16.8 ms observed on Day 1 of the dual-task TDT, lower than any Day 5 thresholds across conditions, and lower than previously reported values. Figure S1 gives an overview of Day 1 thresholds reported in other dual-task TDT studies. Supplementary Table S3 gives the parameters from each of these studies for comparison.

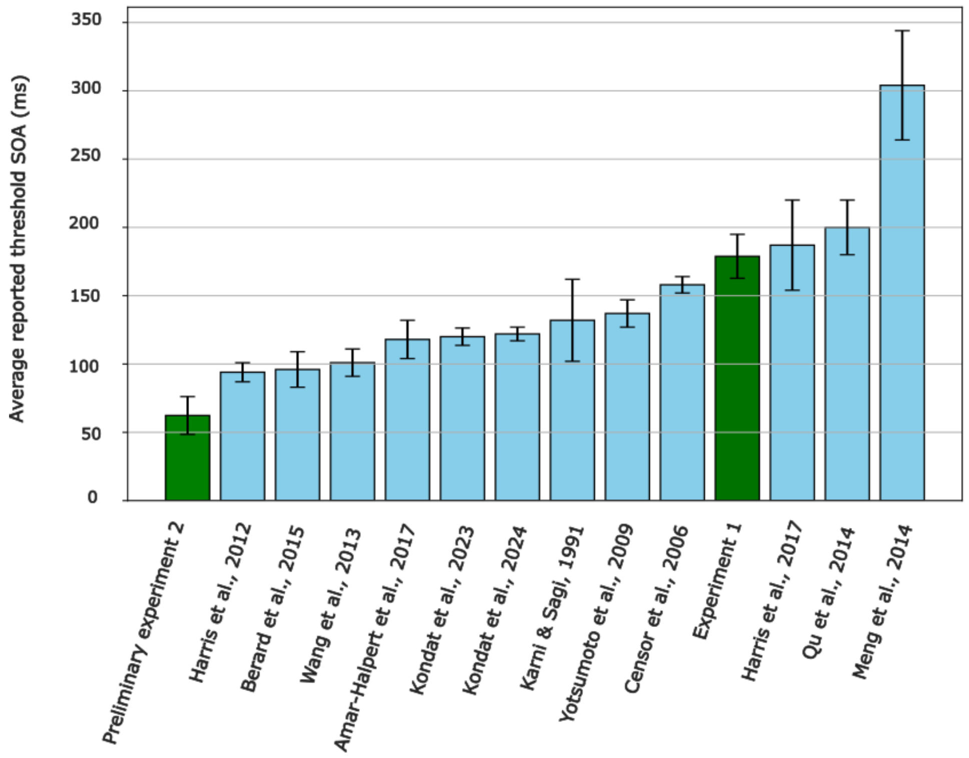


Figure S1: SOA thresholds in a single-task version of the TDT (our data, green bar) compared with SOA thresholds in the standard dual-task version of the TDT in a set of representative studies (light blue bars). Presented numbers are means with SEM, only for Karni & Sagi, 1991 we report the median as no individual thresholds for Day 1 were reported. For parameters used in each study, see Supplementary Table 3.

Table S3: Overview of behavioral paradigms and parameters in related studies, referred to by the first author and publication year (full references in the reference list). Under Method, we refer to random (all SOA levels are presented intermixed) or fixed (blocks of trials are separated per SOA level) stimulus presentation modes. For Censor and Sagi (2008) thresholds, we used the sample that received 26 trials per block (their Fig. 2). For Harris and Sagi (2018) we used the 4-degree eccentricity condition (their Fig. 2). For (Meng et al., 2014) we used only the non-dyslexic participants (children). Some values were extracted using PlotDigitizer (https://plotdigitizer.com), so numbers are approximate.

| **Paper** | **Sample** | **Target** | **Method** | **Eccentricity** | **Trials/soa** | **Trials** | **Feedback** | **Threshold criterion** |
|---|---|---|---|---|---|---|---|---|
| Dual-task TDT | 30 | 17 | Random | $5.72°$ | 18 | 252 | Fixation | 81.6 |
| Preliminary experiment | 10 | 17 | Random | $5.72°$ | 18 | 252 | No | 81.6 |
| Amar-Halpert, 2017 | 24 | 10 | Random | $5.72°$ | 18 | 252 | Fixation | 81.6 |
| Berard, 2015 | 9 | 13 | Fixed | $5--9°$ | 39 | 273 | Fixation | 75 |
| Yotsumoto, 2009 | 6 | 13 | Fixed | $5--9°$ | 26 | 273 | Fixation | 75 |
| Censor, 2006 | 7 | 40 | Fixed | $4.46--6°$ | 26 | 104 | Fixation | 81.6 |
| Karni, 1991 | 6 | 10 | Fixed | $2.5--5°$ | 50 | $\sim$1000 | Fixation | 80 |
| Harris, 2012 | 55 | 10 | Random | $5.3°$ | 18 | 252 | Not reported | Not reported |
| Harris, 2018 | 5 | 10 | Random | $4°$ | 18 | 252 | Fixation | Not reported |
| Kondat, 2023 | 62 | 10 | Random | $5.46°$ | 18 | 252 | Fixation | 81.6 |
| Kondat, 2024 | 41 | 10 | Random | $5.46°$ | 18 | 252 | Fixation | 81.6 |
| Meng, 2014 | 18 | 36 | Staircase | $2.5--5°$ | n/a | 180 | No | 66.7 |
| Qu, 2014 | 24 | 17 | Fixed | $13--19°$ | 16 | 144 | No | 81.6 |
| Wang, 2013 | 8 | 13 | Staircase | $4--6.5°$ | n/a | 80 | Fixation | 79.4 |

**Mixed Linear Model results**

Table S4: Mixed linear model (single-task TDT) with the interaction effect. Reference condition = Control, baseline = Day 1.

| Predictor | $\beta$ | SE | $z$ | $p$ |
|---|---|---|---|---|
| Intercept | 123.675 | 17.294 | 7.151 | $< .001$ |
| Time | -35.983 | 14.539 | -2.475 | .013 |
| condition 2 | -4.730 | 24.458 | 0.193 | .847 |
| condition 3 | 2.683 | 24.458 | 0.110 | .913 |
| Time $\times$ condition 2 | -13.061 | 20.561 | -0.635 | .525 |
| Time $\times$ condition 3 | -67.83 | 20.561 | -0.397 | .691 |

Table S5: Mixed linear model (single-task TDT) without the interaction effect. Reference condition = Control, baseline = Day 1.

| Predictor | $\beta$ | SE | $z$ | $p$ |
|---|---|---|---|---|
| Intercept | 130.061 | 13.466 | 9.659 | $< .001$ |
| Time | -43.060 | 8.161 | -5.276 | <.001 |
| condition 2 | -7.056 | 15.949 | -0.442 | .658 |
| condition 3 | -4.689 | 15.940 | -0.294 | .769 |

To examine whether individual differences in reminder performance were related to overall learning, we tested for a correlation between participants' performance during the reminder sessions (defined as the percentage correct on all the reminder trials) and their overall learning (Day 5 - Day 1). No significant correlation was observed (r = -0.238, p = 0.5072), suggesting that splitting the *reminder* condition based on reminder strength was not warranted.

In addition, to confirm that performance improvements in the *full training* condition reflected genuine learning rather than a test–retest effect, we compared thresholds between Day 2 and Day 5 within this condition. Participants improved significantly from Day 2 to Day 5 (t(9)=4.36, p=.0018), indicating that any results observed where not the results of a simple test-retest effect.

**Tranfser results single-task TDT**

In the single TDT, we tested location transfer in only 11 out of 30 participants, due to time constraints on Day 5. Of those 11, 2 did not reach an 81.6% threshold on the transfer location. Transfer was tested only in three participants of the *full training* condition, two participants of the *reminder* condition, and four from the *control* condition, and as a result no condition differences were inspected. Thresholds were significantly higher at the secondary location compared to the initial location (t(8)=-2.32, p=0.0487, cohen's d=-0.77), indicating specificity.
